# Shared Component Analysis

**DOI:** 10.1101/2020.07.23.218560

**Authors:** Alain de Cheveigné

## Abstract

This paper proposes Shared Component Analysis (SCA) as an alternative to Principal Component Analysis (PCA) for the purpose of dimensionality reduction of neuroimaging data. The trend towards larger numbers of recording sensors, pixels or voxels leads to richer data, with finer spatial resolution, but it also inflates the cost of storage and computation and the risk of overfitting. PCA can be used to select a subset of orthogonal components that explain a large fraction of variance in the data. This implicitly equates variance with relevance, and for neuroimaging data such as electroencephalography (EEG) or magnetoencephalography (MEG) that assumption may be inappropriate if (latent) sources of interest are weak relative to competing sources. SCA instead assumes that components that contribute to observable signals on multiple sensors are of likely interest, as may be the case for deep sources within the brain as a result of current spread. In SCA, steps of normalization and PCA are applied iteratively, linearly transforming the data such that components more widely shared across channels appear first in the component series. The paper explains the motivation, defines the algorithm, evaluates the outcome, and sketches a wider strategy for dimensionality reduction of which this algorithm is an example. SCA is intended as a plug-in replacement for PCA for the purpose of dimensionality reduction.

## 1 Introduction

Electrophysiological and brain imaging data consist of a set of time series of values, one series for each sensor or pixel. Progress in recording techniques has led to more channels, and thus richer data, at the cost of a greater computation and storage burden, and a greater risk of overfitting in analysis or classification. More sensors allow better spatial resolution for source localization, and more degrees of freedom to factor desired and undesired activity into distinct subspaces via spatial filtering. However, this advantage may be offset by overfitting in the data-driven algorithms that are needed to find those spatial filters. Overfitting is an aspect of the “curse of dimensionality”, a prominent concern in pattern recognition and machine learning.

This has led researchers to question the usefulness of higher-density sensor arrays (e.g. Alotaiby et al. 2015; Montoya-Martinez et al. 2019), or to propose discarding channels to avoid overfitting. Given the cost entailed in providing those extra channels in the first place, and the fact that the additional information they provide *should* logically be beneficial, if only to reduce noise, those ideas feel unsatisfactory. There must be better ways to make full advantage of the highdimensional data, and that is what this paper is about.

We are often interested only in a small number of brain sources (e.g. their time course or spatial extent), that are linearly mixed with a larger number of nuisance sources. Despite an unfavorable signal-to-noise ratio (SNR) at each sensor, it may be possible to recover target sources, in some cases perfectly, by applying appropriately-designed spatial filters. More sensors allow the filters to be more effective, and enhances the spatial detail of the recovered sources. Unfortunately, in the absence of precise anatomical information concerning the geometry and properties of the sources and conductive medium, the only way to find those filters is via data-driven algorithms. Those algorithms are prone to overfitting, and that tendency is worse if the data have many channels. We may then find ourselves in the uncomfortable situation that adding more observations actually *degrades* the outcome, as schematized in Fig. 1.

**Figure 1:**
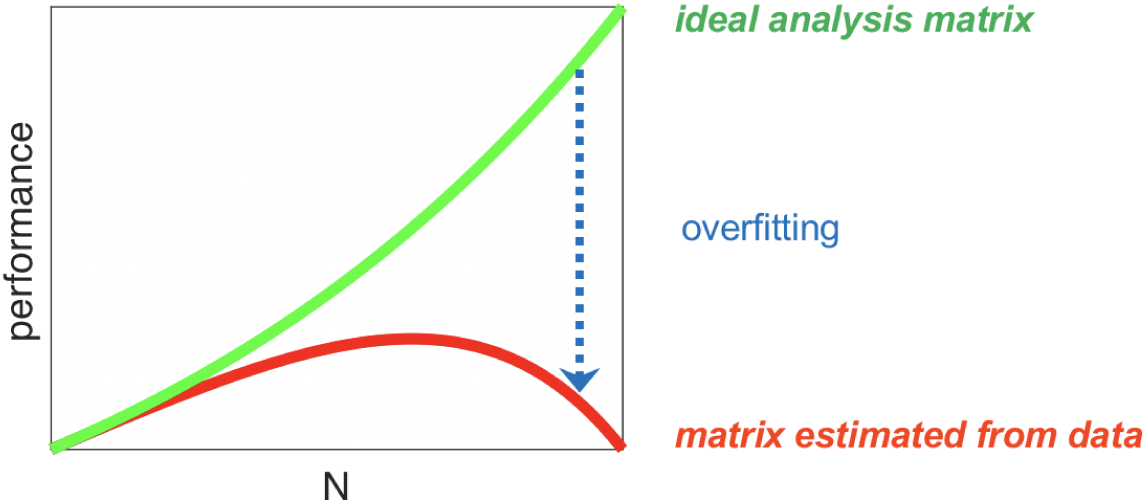
The curse of dimensionality. Increasing the number of channels is expected to boost the performance of methods that rely on linear analysis to extract a source from noise (green), but this may be offset by overfitting, causing performance to peak at some intermediate number of dimensions and decline beyond (red).

A straightforward solution is to discard channels, so as to move back along the tradeoff and reach the peak (Fig. 1). This may be beneficial, but it is unsatisfying because it implies discarding observations. Slightly more sophisticated is the widespread practice of applying *Principal Component Analysis* (PCA) and discarding principal components (PCs) of high rank and low variance. This procedure implicitly equates relevance with variance, which makes sense if we consider that instrumental noise (for example produced by each sensor) induces a “floor” of uncorrelated variance that would swamp any dimension with variance below that floor. Above that floor, however, interesting brain activity might have less variance than noise and artifact, and excluding dimensions on the basis of variance runs the risk of missing it.

There exists a very general procedure that allows variance differences to be neutralized and replaced with some other criterion. It involves three steps: (a) whiten the data by applying PCA and then normalize, (b) apply a “bias filter” to emphasize directions that maximize the criterion of interest, (c) apply a second PCA to rotate the data and reveal those directions. This procedure, which underlies many analysis methods (Parra et al., 2005; Särelä and Valpola, 2005; de Cheveigné and Parra, 2014), can be used to isolate one or a small set of optimal components for further analysis or interpretation. We can also consider it more conservatively as a means to reduce dimensionality, by discarding components that score poorly on the criterion.

A weakness of this procedure is that the initial step (a) confers equal weight to all directions in the data, including those of weak variance likely to reflect noise. If some combination of those dimensions (often numerous) happens to score high on the criterion, by chance, it may end up being selected in step (c), an example of overfitting. This can be addressed by truncating the series of PCs after step (a), but then we are then back at square one: variance determines which directions are retained, which is what we set out to avoid.

This paper investigates an alternative to PCA based on the assumption that desired sources are likely to impinge on *multiple sensors*. For example, sources within the brain may be observed on multiple electrodes in electroencephalography (EEG), or sensors in magnetoencephalography (MEG), or pixels or voxels of an imaging technique. This method, dubbed SCA, is designed to serve as a “plug-in replacement” for PCA.

## Methods

This section describes the algorithm, and provides full details of the evaluation methods and experiments. The busy reader is encouraged to read the sections on Conceptual Tools and SCA Algorithm, then skip to Results and return for details as needed.

### 1.1 Conceptual tools

#### Data Model

Data consist of a matrix **X** of dimensions *T* (time) *× J* (measurement channels). All channels are assumed to have zero mean. Each channel is a weighted sum of sources, desired (e.g. within the brain) or undesired (e.g. artifact):

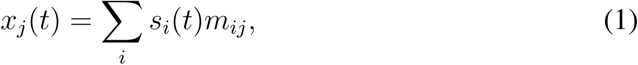

where *t* is time, [*s*_*i*_(*t*)], *i* = 1 … *I* are sources, and the *m*_*ij*_ are unknown sourceto-sensor mixing weights. In matrix notation **X**=**SM**. This model matches the physical mixing process which is, to a good approximation, linear and instantaneous.

#### Linear analysis

Sources of interest may have low SNR at every electrode or sensor, in which case our best option to retrieve them is to combine information across channels with weights *u*_*jk*_:

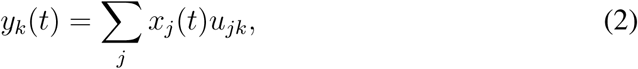

where each “component” *y*_*k*_(*t*) is a weighted sum of the observations. In matrix notation **Y**=**XU**, where **Y** is a matrix of several such components and **U** is the analysis matrix. Each column of **U** defines a *spatial filter* that can be used to enhance target sources and suppress noise. Ideally, **U** should be the inverse of **M** (i.e. an “unmixing matrix”). Unfortunately, that inverse usually does not exist, if only because there are many more sources than observations. At best we hope that some of the *y*_*k*_(*t*) are less affected by noise than are the raw observations *x*_*j*_(*t*).

The analysis matrix **U** can be predetermined (spatial smoothing, gradient, Laplacian, rereferencing, etc.) or else derived from the data. A wide range of data-driven techniques are available, such as PCA, ICA (Independent Component Analysis, Hyvärinen et al. 2009), LDA (Linear Discriminant Analysis), beamforming, Sekihara et al. (2006), CCA (Canonical Correlation Analysis, Hotelling 1936), CSP (Common Spatial Patterns, Koles et al. 1990), regression, and so-on. Two phases should be distinguished: *training*, in which the model is fit to the data, yielding the analysis matrix **U**, and *testing* in which that matrix is applied. These phases are confounded if the matrix is trained and applied to the same data set (as in common parlance “PCA was applied to the data”). Training instead on one dataset and testing on another (cross-validation) allows to control for *overfitting*, which occurs when a model fits details of the training set that do not generalize to new data.

#### Vector spaces

Given our interest in weighted sums, it is useful to reason in terms of *vector spaces* (which are complete for that operation). Each time series can be represented as a “point” within such a space. The *I* sources *s*_*i*_(*t*) span a space that contains all their linear combinations, among them the observed signals *x*_*j*_(*t*) (Eq. 1) that themselves span a smaller *subspace* that contains all their linear combinations, among them the linear analysis components *y*_*k*_(*t*) (Eq. 2).

Vector spaces provide useful concepts such as dimensionality, projection, orthogonality etc. The *dimensionality* of the source space is determined by the number of linearly independent sources (brain or other) which is potentially very large.

That of the subspace spanned by the electrodes/sensors/pixels is much smaller, *J* or less. In the measuring process, brain and noise source signals are all *projected* onto that subspace, and we know them only by these projections. A goal of linear analysis is to further project those observations onto a subspace devoid of noise. Conveniently, some of these methods (e.g. PCA) produce a basis of signals that are mutually *orthogonal*, i.e. pairwise uncorrelated.

#### Curse (and blessing) of dimensionality for data-driven analysis

Signal and noise are more likely to be separable if the sensor data space has more dimensions. Indeed, to find an analysis matrix **U** that projects to a clean signal subspace, the complement of that space (the noise subspace) must be large enough to fully span the noise. That is more likely to be true if there are many sensors/electrodes/pixels/voxels to start with: the more channels the better. Such is the “blessing” of dimensionality.

The counterpart of large dimensionality is a greater risk overfitting, limiting the effectiveness of data-driven methods to find **U**. This is the *curse* of dimensionality: we may well be in the annoying position that an effective demixing matrix **U** exists, but we are unable to find it. The purpose of *dimensionality reduction* or *regularization* (closely related, Tibshirani et al. 2017) is to mitigate this problem. Dimensionality reduction helps by reducing the number of free parameters in the analysis model.

A trivial form of dimensionality reduction is to simply discard channels, or eschew opportunities to increase their number. This is justified if there is good reason to believe that those extra channels add no helpful information, otherwise it is wasteful in resources and information. A more sophisticated strategy is to apply PCA and retain only the high-variance PCs. This justified if low-variance dimensions are uninformative, for example swamped by a noise floor; if not PCA can actually be harmful, and the same might apply to regularization methods such as ridge regression that are closely related to PCA (Tibshirani et al., 2017). This justifies considering alternatives to PCA.

An interesting parallel can be made with the computer science concept of *search*: data-driven methods such as regression or ICA can be understood as methods to search for the optimal analysis matrix **U**. Dimensionality reduction, then, is akin to *pruning* the search space.

### 1.2 The SCA algorithm

We wish to replace variance with a plausible property of the mixing matrix **M**. Sources sufficiently distant from sensors (e.g. within the brain) are likely to impinge on multiple EEG or MEG channels, whereas sources closer to the sensors (e.g. sensor noise or myogenic artifact) are likely to each impinge each on one or a few channels. This assumption, which equates relevance with “sharedness” across channels, does not need to be strictly verified for the data at hand. It needs only to do a better job than variance for the purpose of favoring relevant dimensions. The principle of the algorithm is simple: normalize each column of the observation matrix **X** so as to neutralize any between-channel variance differences, apply PCA, and select the first PC representing the direction of highest variance after PCA rotation. That direction is projected out of the measurement matrix, and the operation repeated, as described below.

#### The algorithm

Starting with the *J* -column matrix **X** of mean-removed data, and setting *j* → 1, **X**^*’*^ → **X**, the following steps are performed to obtain the component matrix **Y**:

1. Normalize each column of **X**^*’*^,
2. Apply PCA, save the first PC as *y*_*j*_(*t*),
3. Project *y*_*j*_(*t*) out of **X**^*’*^,
4. *j* ← *j* + 1, go to step 1 until *j* = *J*.

The algorithm can be stopped at *J*^*’*^ *< J* if we need fewer components than channels. If a column of **X**^*’*^ is all zeros (i.e. its norm is zero) step 1 is omitted to avoid division by zero. The columns of **Y** produced are mutually uncorrelated (or zero). The rationale is the following. If all columns of **X** were mutually uncorrelated (no shared variance), all PCs in step 2 would have variance one. Thus, a PC with variance greater than one implies the existence of components shared between channels. The first PC reflects the one most shared: projecting it out of the data (step 3) and renormalizing (step 1 of the next iteration) allows the next most-shared component to be isolated, and so-on. Renormalization is necessary because variance might no longer be equal across channels once a component is removed.

#### SCA-based dimensionality reduction

Similar to PCA-based dimensionality reduction, a subset of SCs is retained and the rest discarded. SCA thus serves as a “plug-in replacement” for PCA, retaining dimensions most shared rather than those with greatest variance.

#### Implementation

To save computation, the algorithm operates on the data covariance matrix rather than the data themselves, producing the analysis matrix **U**. The computation cost is dominated by eigenvalue decomposition in step 2, *𝒪*(*J* ^3^) or better, yielding a cost of *𝒪*(*J* ^4^), or *𝒪*(*J*^*’*^*J* ^3^) if *J*^*’*^ components are calculated. A Matlab implementation is provided by the function nt_sca.m of the NoiseTools toolbox (http://audition.ens.fr/adc/NoiseTools/).

### 1.3 Evaluation

#### Task

We consider a task in which a single probe signal is projected onto a multichannel matrix of observations, with the purpose of deciding whether that signal might reflect a “source” within the data. A regression model is fit to the data, and that fit is evaluated by measuring the empirical correlation between signal and projection. Fit (train) and evaluation (test) are performed either on the same data (no cross-validation) or on different data (cross-validation). A large correlation value suggests that the probe reflects a source within the data, but this conclusion might be affected by overfitting.

#### Leave-one-out cross-validation

The probe signal and data are both divided into *N* segments. For each segment *n* = 1 … *N*, a model is fit on the concatenation of the *N* −1 segments complementary to that segment and applied to segment *n*, projections of all segments are concatenated, and the correlation is calculated between concatenated signal and concatenated projections. This provides the estimate of “cross-validated correlation”. Note that, for data with long serial correlation (low-pass spectrum), training and test segments should not be contiguous. Simple correlation (not cross-validated) is obtained by fitting the model and testing it on the same data.

#### Measuring the effect of dimensionality reduction

Overfitting will typically reduce cross-validated correlations. This happens if the model fits too closely to training data points that do not well represent held out data. The benefit of dimensionality reduction can thus be measured as a boost to cross-validated correlation: the larger this boost, the better.

#### Simulations with synthetic data

Four simulations were carried out using three synthetic datasets (*A* to *C*) to better understand the problem addressed and illustrate the method. The first simulation illustrates spurious correlation and its dependency on dimensionality, number of samples, and spectral content, using *dataset A*. The second illustrates the effect of overfitting on the empirical correlation between a known signal vector and a data matrix in which that signal is mixed, using *dataset B*. The third compares PCA and SCA, also using *dataset B*. The fourth also compares PCA and SCA, using *dataset C*.

*Dataset A* consisted of a “probe” signal matrix of size *T ×* 1, with *T* taking values from 1000 to 10000 samples, and a “data” matrix of size *T × J, J* = 1024. Probe and data were unrelated, i.e. their samples were taken from uncorrelated Gaussian distributions. The data matrix was submitted to PCA and subsets of *N*_*pcs*_ PCs were selected, with *N*_*pcs*_ varied as a parameter. The probe was then projected on the selected PCs, and the empirical correlation between the probe vector and the projection vector was estimated, with and without cross-validation (for finite *T*, the empirical correlation is typically non-zero). In one condition, probe and data were additionally low-pass filtered with a Butterworth filter of order two and cutoff 10 or 1 Hz (assuming a sampling rate of 500 Hz), to illustrate the effect of serial correlation.

*Dataset B* consisted of a data matrix of size *T × J, T* = 5000, *J* = 1024, obtained by mixing a matrix of “noise” sources of size *T × J*, via a mixing matrix *M*_*n*_ of size *J × J*, together with a “signal” vector of size *T ×* 1 via a mixing matrix *M*_*s*_ of size 1*×J*. The noise source matrix consisted of Gaussian-distributed values, spatially whitened by PCA followed by normalization so as to have full control of the empirical variance (one for each source) and covariance (0 between pair of sources). The signal consisted of a sinusoid scaled for unit variance. The noise mixing matrix was diagonal with values randomly distributed between 0 and 1, while coefficients of the signal mixing matrix *M*_*s*_ were set to a common gain factor (varied as a parameter), or else to zero. Depending on the condition (as specified in the Results section), signal mixing coefficients were set to zero for channels with noise variance above a threshold, or below a threshold. Thus, the signal could be present only in directions of high variance in the data, or else only in directions of low variance. The data matrix was submitted to PCA (resp. SCA) and the empirical correlation between the signal and subsets of *N*_*pcs*_ PCs (resp. *N*_*scs*_ SCs) was estimated.

*Dataset C* consisted of a “signal” source matrix of size *T × I, T* = 10000, *I* = 128, consisting of sinusoids with different frequencies (thus mutually orthogonal), mixed into a data matrix of same size via a specially-crafted mixing matrix. The first source was mixed into all *J* channels with amplitude 1, the second two sources were mixed into randomly-chosen subsets of *J/*2 channels also with amplitude 1, the next four sources into subsets of *J/*4 channels, and so-on. The last 64 sources were each mixed into one channel. The number of sources per channel depended on the random assignment of sources to channels; here, it ranged from 0 to 11. Uncorrelated Gaussian noise was added to all channels, also with amplitude one, so the data were full rank. All channels were normalized to unit variance and then multiplied by a normally-distributed random number, to ensure that variance is a non-informative cue. The data matrix was submitted to PCA (resp. SCA) and the empirical correlation between selected signals and subsets of *N*_*pcs*_ PCs (resp. *N*_*scs*_ SCs) was estimated.

#### Real data

Two datasets are used, both from a study that aimed to characterize cortical responses to speech for both normal-hearing and hearing-impaired listeners. Sixty four channel EEG responses to speech and tones were recorded in 44 subjects (Fuglsang et al., 2020). Full details can be found in that paper, and the data are available from http://doi.org/10.5281/zenodo.3618205. The EEG data were preprocessed by the Zapline algorithm to remove power line artifacts (de Cheveigné, 2019), high-pass filtered at 0.5 Hz using an order-2 Butterworth filter, and lowpass filtered at 30 Hz using an order-2 Butterworth filter. To remove eyeblink artifacts, a temporal mask was derived from the absolute power on a combination of two EOG channels and three frontal channels (F1, F2, Fz), and the eyeblink components were removed as described by (de Cheveigné and Parra, 2014). We apply PCA and SCA prior to applying two data-driven algorithms (DSS and CCA, as defined below), to evaluate and compare their ability to avoid overfitting.

*Dataset D*, from that study, consisted of EEG in response to a sequence of *M* =180 tones (tone-evoked response). Those data were cut into 180 trials, each starting 1s before and ending 2s after tone onset. Each trial was individually detrended by subtracting a 1st-order polynomial robustly fit to the first and last second of the trial (de Cheveigné and Arzounian, 2018). The first and last seconds worth of each trial were then discarded, leaving a one-second interval post onset for each trial. Outlier trials (contaminated by glitches) were removed based on their Euclidean distance from the average over trials.

The trial matrix was processed by the DSS (denoising source separation) algorithm (de Cheveigné and Parra, 2014) to obtain a transform (spatial filter) to maximize *repeatability* across trials, thus maximizing the SNR of our observation of the brain response (based on the assumption that this response is the same on every trial). The benefit of DSS is quantified by the squared Euclidean distance *d*^2^ between each trial and the average over trials, both normalized to unit variance. This measure ranges from 0 (all trials identical) to 2 (uncorrelated signals, i.e. no repetition across trials): the smaller the value of *d*^2^, the better.

The DSS algorithm is data-dependent and, like regression, prone to overfitting. To *control for* overfitting, the DSS model was fit to subsets of *M* − 1 trials, the optimal spatial filter was applied to the *M* th trial, and *d*^2^ was calculated between that trial and the average over the *M* − 1 other trials. This was repeated for all *M* permutations, and the resulting estimates were averaged. To *reduce* overfitting, prior to DSS the data were submitted to PCA (resp. SCA) and DSS was applied to a subset of *N* PCs (resp. SCs). The relative merits of PCA and SCA are assessed by comparing the minima of *d*^2^ across *N*: the lower, the better.

*Dataset E* consisted of EEG in response to *M* =22 segments of continuous speech, each of 50’ duration. In addition to data in response to a single talker, the database included also data for two talkers (“concurrent speech”), not used here. The EEG data are provided together with the temporal envelope of the speech stimulus, calculated by a model of instantaneous loudness. The speech temporal envelope has been shown to be a predictor of cortical responses (Lalor et al., 2009; Ding and Simon, 2012; Di Liberto et al., 2015; Crosse et al., 2016). The EEG data were cut into 22 trials of 50 s duration, each individually detrended by subtracting a 2nd-order polynomial robustly fit to each channel (de Cheveigné and Arzounian, 2018). The stimulus envelope was filtered by the same combination of high-pass and low-pass filters as the EEG (to avoid introducing a convolutional mismatch). A linear stimulus-response model was fit to the data by projecting the envelope on the EEG (so-called “backward model”). The quality of fit was quantified as the correlation coefficient between stimulus and projection. To compensate for the EEG response latency (unknown and subject-dependent), this process was repeated for a range of time shifts and the shift that produced the largest correlation was retained. Projection and correlation were performed on 30s segments of data, independently for each listener.

This process is prone to overfitting, due to the relatively large number of free parameters (64), limited data duration, and low-pass spectral characteristics of the data. To control for overfitting, correlation was estimated using crossvalidation. To alleviate overfitting, the stimulus was projected on a subset of *N* PCs (resp. SCs) to reduce dimensionality. The relative merits of PCA and SCA are assessed by comparing the maxima of correlation across *N*: the larger, the better.

## Results

We first explore the basic properties of the method using synthetic data, and then move on to real EEG and MEG data to see the benefit extends to practically useful scenarios.

### 1.4 Synthetic data

#### Spurious correlation

The first simulation used *dataset A*. The dataset consists of a “probe” signal vector, and a 1024-channel data matrix consisting of noise drawn from a distribution uncorrelated with the probe. The data matrix was submitted to PCA, and the empirical correlation between the probe and subsets of selected PCs was estimated with and without cross-validation. Figure 2 (left) shows correlation as a function of *N*_*pcs*_ without (thin line) and with (thick line) cross-validation. Without cross-validation, the correlation score approaches 0.5 for large *N*_*pcs*_, despite the fact that the probe and data are drawn from independent processes. This is a classic example of overfitting: more channels or components makes it easier for the fitting algorithm to find some linear combination of the data that is well correlated with the probe. With cross-validation, the correlation estimate is not biased in this way, although it hovers at some distance from zero due to residual (unbiased) correlation. The spurious empirical correlation is larger when there are fewer samples (small *T*) than more (Fig. 2, center). Furthermore, for a given *T*, it is larger if the data are not spectrally white (Fig. 2, right). This is because the serial correlation characteristic of spectrally-colored data reduces the “effective” number of samples in the data.

**Figure 2:**
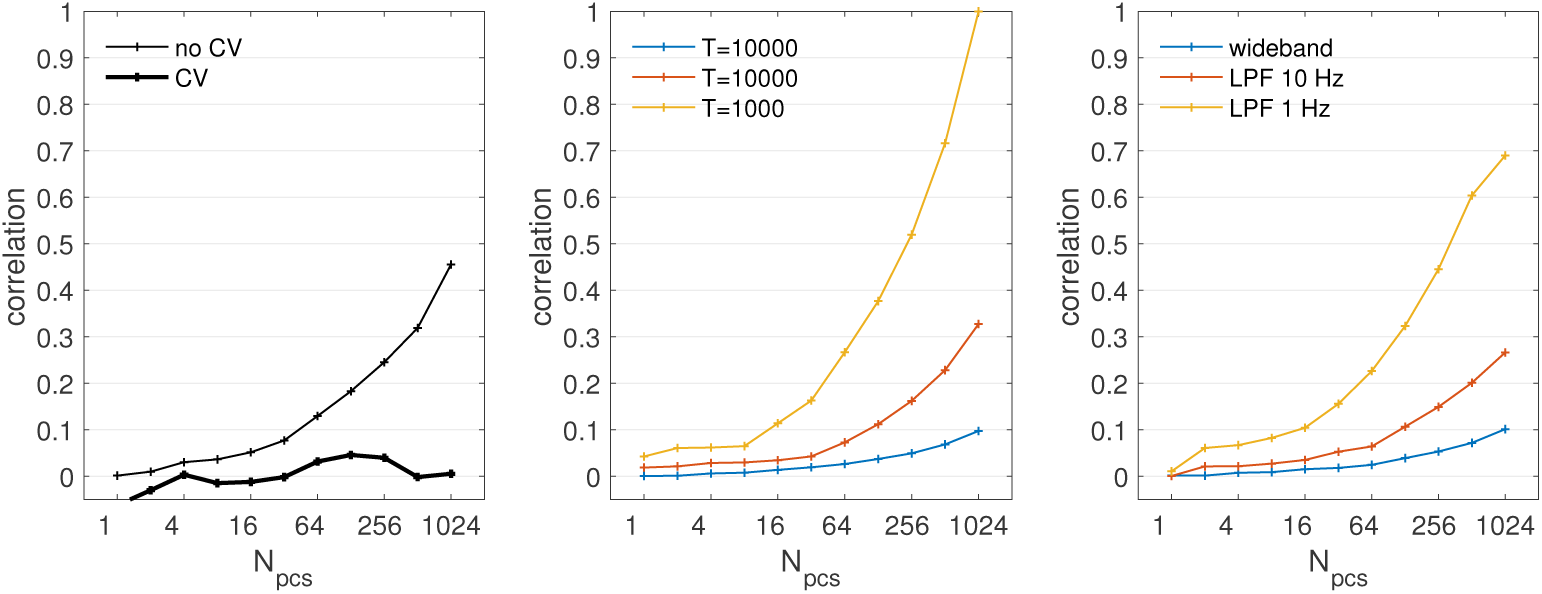
Spurious correlation between a vector of Gaussian noise and its projection on subsets of PCs of a 1024-channel matrix of Gaussian noise nominally uncorrelated with the vector. Left, thin line: empirical correlation estimated without cross-validation, thick line: with cross-validation. Non-zero values reflect spurious correlations. Center: correlation without cross-validation for data matrices with different numbers of rows T. Right: same for T=100000, for data that have undergone different degrees of filtering, including none (wide-band). Spurious correlations are greater for more than for fewer dimensions, for fewer than for more data samples, and for spectrally coloured than for wide-band data.

This simulation illustrates the well-known fact that overfitting increases with the number of dimensions (*N*_*pcs*_), decreases with the number of data samples (*T*), and is exacerbated if the data are not spectrally-white.

#### Regression and overfitting

The second simulation used *dataset B*. That dataset consists of a “signal” vector and a simulated data matrix within which this signal has been mixed together with noise. The purpose of this simulation is to illustrate the fact that some datasets benefit from dimensionality reduction, others not: there is no “one size fits all” strategy. Real data can differ widely in how signal components are distributed over dimensions within the data, in particular whether they coincide with dimensions of low noise variance, or instead high noise variance. To simulate these two situations, the noise mixing coefficients were randomly distributed between 0 and 1 (i.e. some channels had high noise variance, others low), and the signal was mixed with a gain factor that depended on the noise variance.

In Fig. 3 (left), the signal gain factor was 0 for channels in which the noise mixing coefficient was less than 0.7, and 0.02 otherwise. In other words, *the signal was present only within the 70% of channels with high noise variance*. After PCA, the signal was projected on subsets of PCs and the correlation coefficient between signal and projection was estimated with cross-validation (thick line) and without (thin line). Without cross-validation, the correlation grows monotonically with *N*_*pcs*_, with cross-validation it peaks at *N*_*pcs*_=128 and decreases beyond. This non-monotonic pattern is an often-reported result of overfitting (Tibshirani et al., 2017).

**Figure 3:**
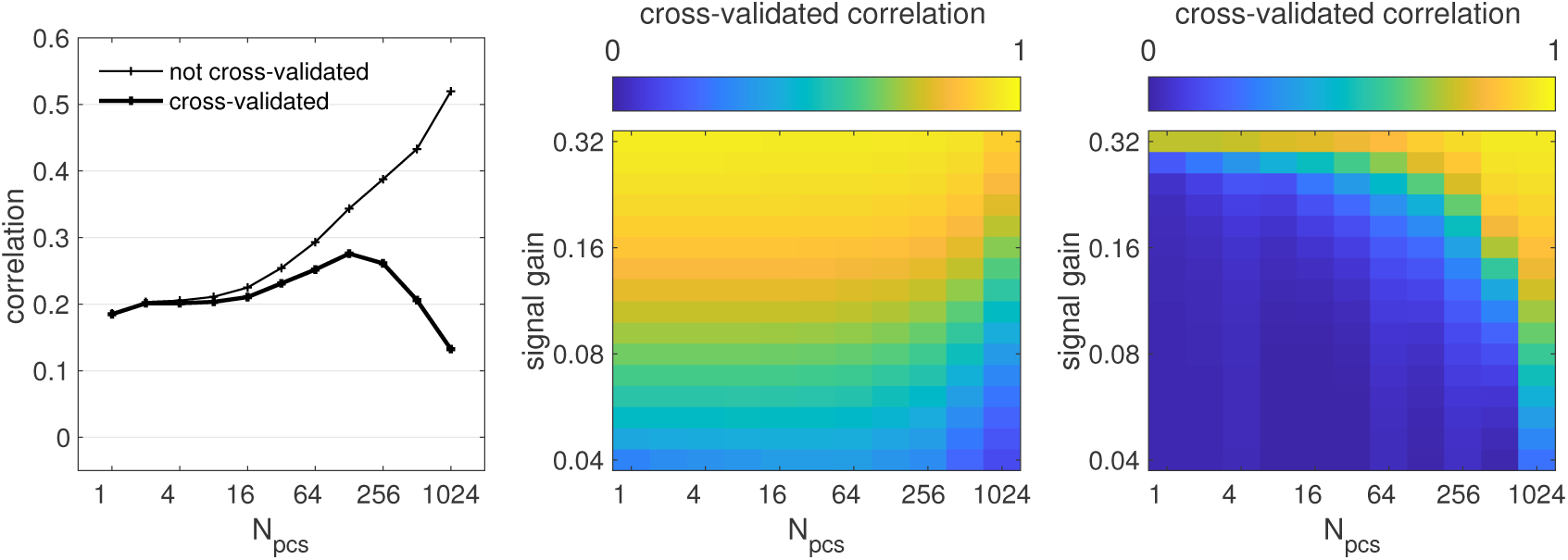
Correlation between a signal vector (a sinusoid) and its projection on selected PCs of a 1024-channel data matrix in which that signal has been mixed together with uncorrelated Gaussian noise. Left, thin line: without cross-validation, thick line: with cross-validation. Cross-validated correlation peaks at an intermediate value of N_pcs_ and decreases beyond. Center: cross-validated correlation as a function of N_pcs_ for different values of SNR, coded on a color scale, for a data set in which the signal was present only on channels of high variance. Cross-validated correlation is greater for small or intermediate N_pcs_. Right: same, but the signal was present only on channels of low variance. Cross-validated correlation is greatest for N_pcs_=1024 (i.e. no dimensionality reduction). The benefit of PCA-based dimensionality reduction depends on the data.

However, this non-monotonic pattern is not always observed. In Fig. 3 (center), the signal gain factor was 0 for channels for which the noise mixing coefficient was less than 0.9, and for the other channels it was set to a value that was varied between 0.02 and 0.32 as a parameter. The value of the cross-validated correlation is plotted on a color scale as a function of *N*_*pcs*_ and gain. For small values of gain, the correlation seems to vary non-monotonically as in Fig. 3 (left). For larger values of gain it decreases monotonically with *N*_*pcs*_. In both cases, it is beneficial to discard low-variance PCs.

In contrast, in Fig. 3 (right), the signal gain factor was 0 for channels for which the noise mixing coefficient was greater than 0.3, and for the other channels it was to a value that was varied between 0.02 and 0.32 as a parameter. In other words, *the signal was present only on channels with low noise variance*. The crossvalidated correlation increases monotonically with *N*_*pcs*_. For these data, it is *not* beneficial to discard low variance PCs.

These gain parameters and cutoffs were chosen arbitrarily for visual clarity, to illustrate the fact that PCA-based dimensionality reduction can be either beneficial or detrimental, depending on whether data directions of high variance coincide with directions of high SNR, or not.

#### SCA can be more effective than PCA

This third simulation also used *dataset B*. Figure 4 compares the values of correlation obtained with PCA (black) and SCA (red) as a function of the number of components retained. Dotted lines are without cross-validation, full lines are with cross-validation, and the two panels are for data with different properties. For the left panel, the 30% of channels with lowest variance carried the signal, with gain=0.02. For the right panel, only the 30% of channels with highest variance carried the signal, with gain=0.1. These parameter values are arbitrary, chosen merely for visual clarity.

**Figure 4:**
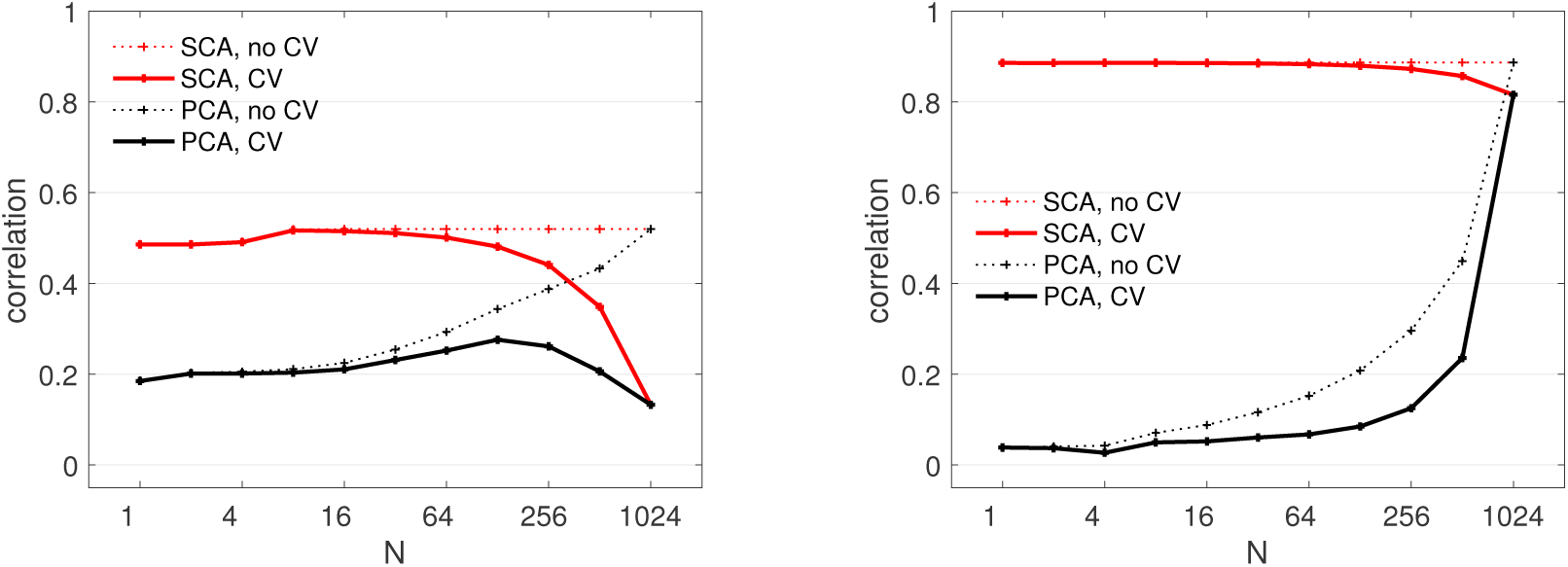
Comparing correlations between probe signal and dimensionalityreduced data for SCA and PCA with and without cross-validation, for synthetic data. Both panels show correlation as a function of the number of PCs (black curves) or SCs (red curves), with cross-validation (full lines) and without cross-validation (dotted lines), for synthetic datasets with different parameters. Left: the signal was present only on the 30% of channels with lowest variance, SNR=0.02. Right: the signal was present only on the 32% of channels with highest variance, SNR=0.1. Cross-validated correlation peaks at an intermediate value of N_pcs_ for one dataset and not the other, while non-cross-validated correlation increases monotonically in every case (dotted lines). On both datasets, SCA performs better (i.e. allows higher cross-validated correlation values) than PCA.

Focusing on cross-validated correlation (full lines), the values for PCA and SCA are equal for the largest number of components, as expected because both sets of 1024 components span the full data. For all other values of *N* the correlation values are greater for SCA than for PCA, suggesting that for these particular data, sharedness is a more useful criterion than variance. Given that we are free to choose *N*, we are primarily interested in the *maximum* of cross-validated correlation across *N*. Importantly, this maximum is greater for SCA than for PCA, and the fact that it is obtained for smaller *N* may also be advantageous. It should be cautioned that these results were obtained with a particular type of data: a single “signal” distributed across a large number of sensors, “noise” uncorrelated across sensors, and variance tailored to be uninformative or misleading (to the chagrin of PCA).

This simulation demonstrates that, for some data, SCA can serve advantageously as a replacement for PCA.

#### SCA works also with multiple shared sources

This simulation used *dataset C*, in which 128 sinusoidal sources are mixed into 128 channels together with Gaussian noise, such that the first source is mixed into all channels, the second two are each mixed into 64 randomly-chosen channels, and subsequent sources into progressively smaller numbers of channels. This mimics EEG or MEG data where some sources have a wider spatial extent than others.

The data were submitted to SCA (resp. PCA), subsets of contiguous SCs (resp. PCs) starting at 1 were selected, individual sources were projected on these subsets, and cross-validated empirical correlation was estimated between source and projection. Figure 5 (right, dotted lines) shows, for the first 12 sources, the correlation between that source and its projection on subsets of PCs (black) or SCs (red). To be precise: the data point at e.g. *N* =50 is the correlation between the source and the subspace spanned by components 1 to 50. Values for source 7 are shown as continuous bold lines. Correlation coefficients increase rapidly for SCA, indicating that the initial sources are highly correlated with subspaces spanned by a small number of initial SCs. For PCA they increase more slowly, suggesting that PCA is less effective in “concentrating” relevant dimensions of the data.

**Figure 5:**
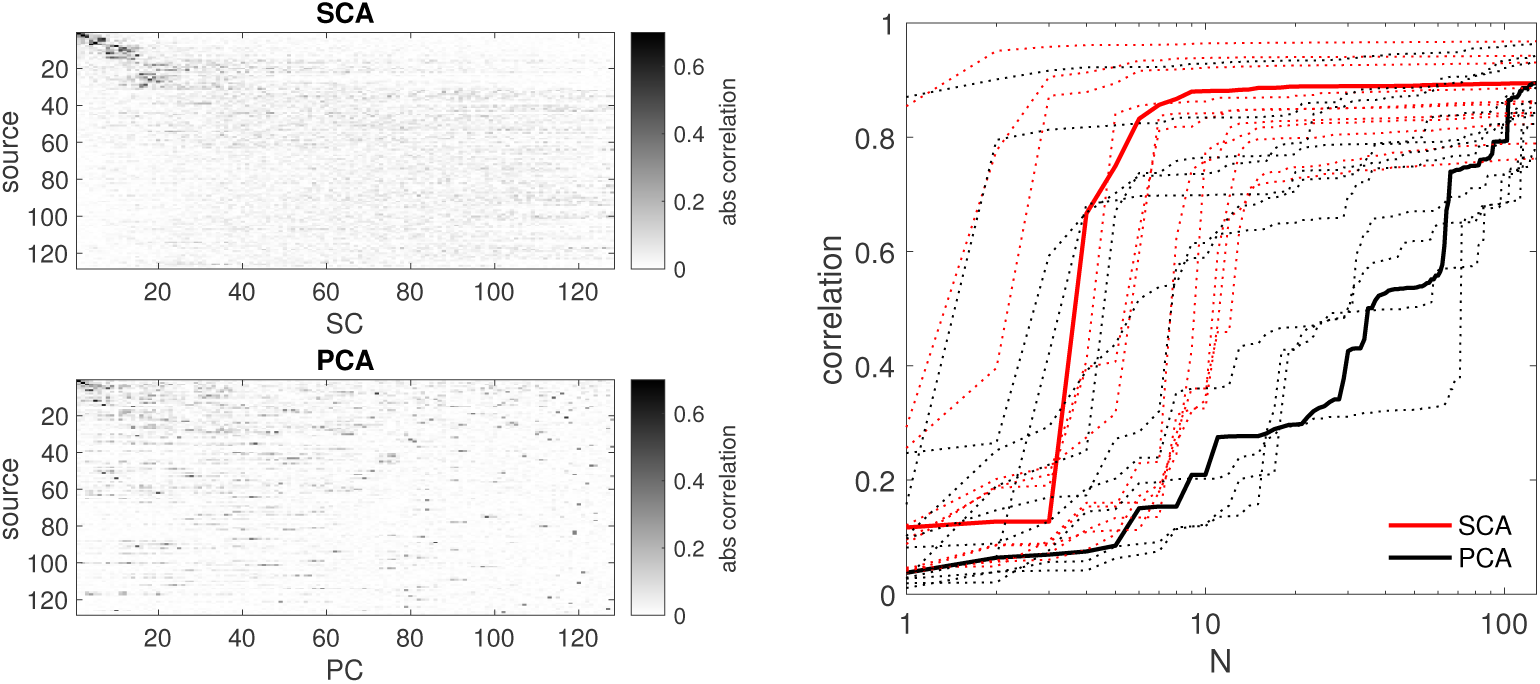
Synthetic data, multiple shared sources. Left: correlation between individual sources and SCs (top) or PCs (bottom). The first few sources (most shared across channels) are more tightly concentrated within the first few SCs than within the first few PCs. Right: correlation between each source and its projection on the first N SCs (red) or PCs (black), for the first 12 source. Continuous bold lines are components for source 7. A steeper increase of correlation with N for SCA implies that it is more effective than PCA in “packing” useful variance within a small number of components.

Figure 5 (left) shows the absolute values of correlation coefficients between individual sources and SCs (top) or PCs (bottom). Correlation coefficients tend to be large between the first sources (most widely shared across channels) and the first SCs, whereas they are smaller between first sources and later SCs, and between first SCs and later sources. This suggests that SCA has indeed managed to “concentrate” the initial sources within the initial SCs. For PCA, the correlation coefficients between sources and PCs are more widely distributed, suggesting that PCA is less effective in gathering the initial, widely-shared, sources.

This simulation shows that the benefit of SCA over PCA can extend to situations where there are several sources superimposed within the data, each shared across sensors. Again, it should be cautioned that this result was obtained with a data set crafted such that variance was relatively uninformative, putting PCA at a disadvantage.

### 1.5 Real data

#### EEG response to tones

This simulation used *dataset D* (EEG responses to 180 repetitions of a tone, measured from 44 subjects, see Methods). PCA (resp. SCA) was applied to the preprocessed data and subsets of *N* PCs (resp. SCs) were selected before applying the DSS algorithm (Methods) to find an optimal spatial filter to enhance repeatability (de Cheveigné and Parra, 2014). The DSS algorithm is effective to suppress the background EEG activity that masks the regularly repeating brain response to each tone (compare Fig. 6 left and center). As a data-driven algorithm, it is prone to overfitting (the algorithm can find particular linear combinations that happen to reoccur by chance over trials), which is why it is worth considering reducing dimensionality before applying it.

**Figure 6:**
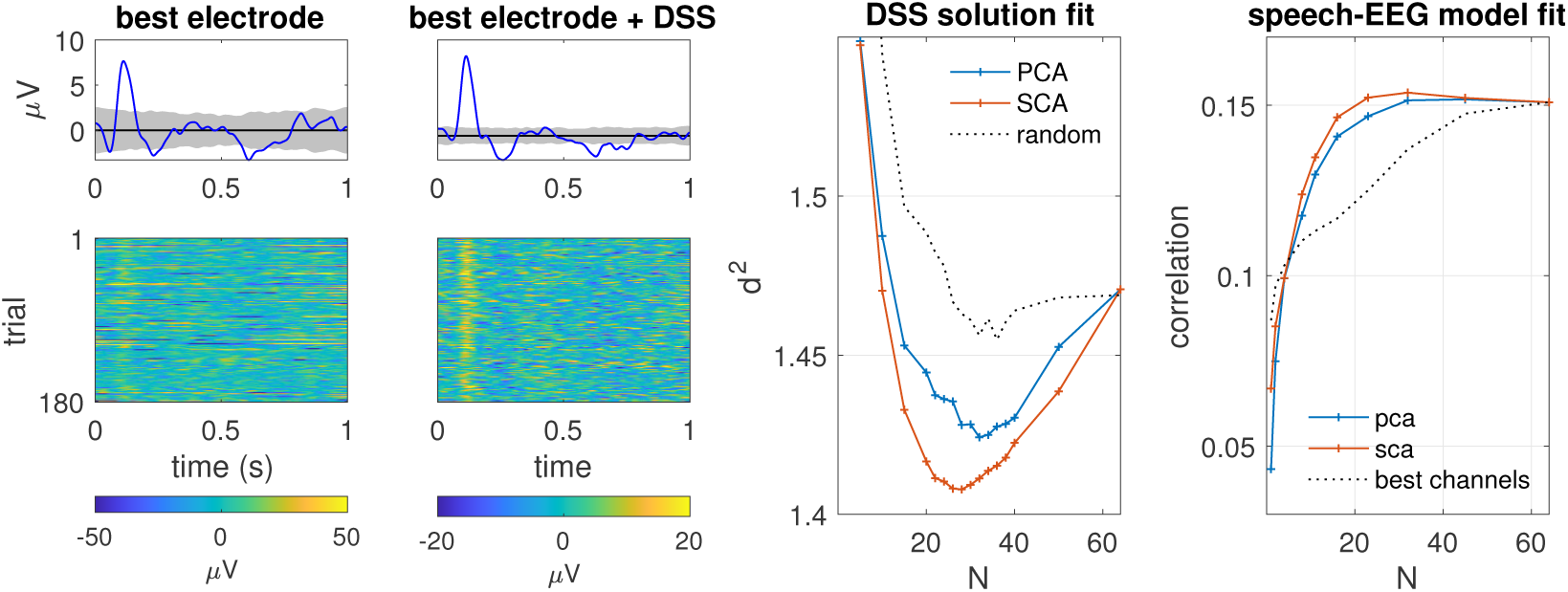
EEG, auditory evoked potentials. Left, top: average over trials (blue) and 2SD of bootstrap resampling (gray) of auditory evoked response for electrode with largest SNR. Next to left: same after denoising with the DSS algorithm. Next to right: quality metric (quantified as d^2^, Methods) as a function of the number of selected PCs (blue) or SCs (red) for one subject. Smaller is better. Dimensionality reduction improves the quality of the DSS solution (dip at at N < N_channels_) and the improvement is greater for SCA than PCA. Right: correlation between speech envelope and projection on a set of N first PCs (blue) or SCs (red). The dotted line shows correlation for a set of N channels, ordered in terms of decreasing correlation with the speech envelope.

Applying PCA and selecting the first *N* PCs is effective in reducing the *d*^2^ metric for left-out trials (Fig. 6, right, blue, compare minimum to value for *N* =64). The best model (lowest *d*^2^) is obtained for *N* ≈ 35. However, applying instead SCA and selecting the first *N* SCs yields an even better model for *N* ≈ 30, on average over subjects. It is worth noting for some subjects PCA performed better than SCA (not shown). Nonetheless a Wilcoxon signed rank test applied to average across *N* for each subject suggests that the benefit of SCA over PCA is reliable at the population level (p=0.004).

The dotted line in Fig. 6 (right) shows the *d*^2^ obtained if the data are transformed by a random matrix (rather than PCA or SCA), and a subset of *N* transformed components is selected. The benefit of this simple form of “dimensionality” reduction, if any, is small, suggesting that any reduction in overfitting is offset by a reduced ability of DSS to resolve the signal within a smaller space. PCA and SCA are both better able to prioritize “good” dimensions, and in this role SCA seems to have an edge over PCA. This example shows that SCA is a viable alternative to PCA in an analysis pipeline for real EEG data.

#### EEG response to speech

This simulation used *dataset E* (EEG responses to 16 segments of speech each of duration 55 seconds from 44 listeners, see Methods). PCA (resp. SCA) was applied to the data and subsets of *N* PCs (resp. SCs) were selected and fit, together with a temporal envelope representation of the stimulus, by a stimulus-response model based on CCA (see Methods) (de Cheveigné et al., 2018). The quality of the model fit was estimated by calculating the correlation between the first pair of canonical correlates (one derived from the stimulus envelope, the other from the EEG) with cross-validation. Specifically: the model was fit to a subset of 15 trials and tested on the 16th (left out), and his operation was repeated with all choices of left-out trial. The correlation values were averaged over folds.

The CCA-based algorithm is data-driven (it finds the linear combination of EEG channels most highly correlated with its projection on the space spanned by time-shifted stimulus envelope signals), and as such it is susceptible to overfitting, that one can hope to counteract by reducing dimensionality using PCA or SCA. Figure 6 (right) shows cross-validated correlation between the first CC pair as a function of the number of components retained, for PCA (blue) and SCA (red), averaged over subjects. SCA produces larger correlation values than PCA, with a peak that occurs at a smaller value of *N*. This pattern is observed for most subjects, albeit not all (not shown), as confirmed by a Wilcoxon signed rank test applied to the average across *N* for each subject (p<10−4). This example also shows that SCA is an alternative to PCA in an analysis pipeline for real EEG data. Also shown in Fig. 6 (right, dotted) is the correlation obtained by projecting the speech envelope on subsets of *electrodes*, selected here based on their individual correlation with the speech envelope (i.e. best first). Interestingly, correlation is considerably lower than for PCA or SCA (both based on the full set of 64 channels), and increases monotonically withe the number of channels up to 64. There is no evidence that reducing the number of channels reduces overfittiing (in contrast to e.g. Alotaiby et al. 2015; Montoya-Martinez et al. 2019; Fuglsang et al. 2017), or that some form of channel selection might be preferable to reducing dimensionality via PCA or SCA.

## Discussion

This paper proposed SCA as a plug-in replacement for PCA for the purpose of dimensionality reduction.

### SCA vs PCA

SCA is similar to PCA in that it decomposes *J* -channel data into *J* mutually-orthogonal components. Similar to PCA, it can be used to implement dimensionality reduction as an early processing stage to reduce overfitting in subsequent data-driven analyses methods. SCA differs from PCA in that it selects components based on *shared variance* rather than variance. It thus gives priority to directions with activity that is spread over multiple channels (sensors, electrodes, pixels, voxels). Such might be the case of brain sources deep enough to impinge on multiple channels, as opposed to more superficial sensor or myogenic noise sources that each impinge on a smaller number of sensors. Reducing dimensionality allows data-driven algorithms to operate within a smaller space, possibly with greater chance of success. We showed several examples, based on synthetic and real data, for which SCA performed better than PCA in this role.

It should be stressed that it is *not* always the case that SCA is better than PCA. The relative value of SCA and PCA depends entirely on the nature of the data, specifically on which of the two criteria (variance or shared variance) better maps to relevance. It is hard to predict which will be the case for arbitrary data. The best strategy may be to try both and choose whichever gives the best results empirically, using appropriate cross-validation and statistical tests to control for double-dipping (Kriegeskorte et al., 2009). The choice (PCA, SCA, or some other method) can be made automatically within a hyperparameter search loop, akin to determining a regularization parameter.

### Multistage dimensionality reduction

Linear data analysis usually involves applying an analysis matrix to the data and selecting one or a few “components”, hopefully more representative of brain processes than the raw data. A very wide range of methods are available, including PCA, ICA, beamforming, CSP, DSS, CCA, and others. Some, such as DSS, can be applied with a wide range of criteria (de Cheveigné and Parra, 2014). It is natural to want to apply them in succession so as to benefit from their cumulated effects. Unfortunately, this does not work: the criterion applied in the last step undoes the effect of any previous transform (as long as that transform is invertible). It is, however, possible to reap benefits of multiple methods if they are used as successive stages of *dimensionality reduction*. Applying, say, an initial stage of PCA and discarding low-variance PCs allows the next stage (e.g. DSS) to work within a smaller space. Discarding DSS components with low scores then allows the next stage (e.g. CCA) to operate in a yet smaller space, and so on. Multistage dimensionality reduction is a plausible way to reap the benefits of multiple criteria of how data should look. SCA adds to this panoply of methods.

### Dimensionality reduction and regularization

It is customary to combat overfitting by applying *regularization*, for example ridge regularization in which some small value is added to the diagonal of the data covariance matrix. It has been pointed out that there is a close connection between ridge regularization and PCA-based dimensionality reduction (Tibshirani et al., 2017). Intuitively, adding a constant to the diagonal of the covariance matrix has the same effect as adding to each data channel a noise signal uncorrelated with the data, and uncorrelated with other channels, which has the effect of “swamping” dimensions of low variance so that they do not affect the solution. This is akin to discarding them, as with PCA. Empirically, there is often not a marked difference between different regularization methods (Wong et al., 2018), and thus no strong basis to prefer any of them over mere dimensionality reduction, which allows variance to be replaced by other criteria (as with SCA) and opens the perspective of multistage linear dimensionality reduction (previous paragraph) and possibly other more sophisticated non-linear (manifold) dimensionality reduction schemes (Burges, 2009).

## Conclusion

This paper investigated Shared Component Analysis (SCA) as an alternative to Principal Component Analysis (PCA) for the purpose of dimensionality reduction of neuroimaging data. Whereas PCA selects orthogonal components that explain a large fraction of variance in the data, SCA instead selects components that contribute to observable signals on multiple sensor channels. Depending on the data, this can result in more effective dimensionality reduction, as illustrated with synthetic and real EEG data. SCA can be used as a plug-in replacement for PCA for the purpose of dimensionality reduction.

## Acknowledgements

Malcolm Slaney and Søren Fuglsang provided useful comments on early drafts of this paper. This work was supported by grants ANR-10-LABX-0087 IEC, ANR-10-IDEX-0001-02 PSL, and ANR-17-EURE-0017.

